# VIQoR: a web service for Visually supervised protein Inference and protein Quantification

**DOI:** 10.1101/2021.06.01.446512

**Authors:** Vasileios Tsiamis, Veit Schwämmle

## Abstract

**Motivation:** In quantitative bottom-up mass spectrometry (MS)-based proteomics the reliable estimation of protein concentration changes from peptide quantifications between different biological samples is essential. This estimation is not a single task but comprises the two processes of protein inference and protein abundance summarization. Furthermore, due to the high complexity of proteomics data and associated uncertainty about the performance of these processes, there is a demand for comprehensive visualization methods able to integrate protein with peptide quantitative data including their post-translational modifications. Hence, there is a lack of a suitable tool that provides post-identification quantitative analysis of proteins with simultaneous interactive visualization.

**Results:** In this article, we present VIQoR, a user-friendly web service that accepts peptide quantitative data of both labeled and label-free experiments and accomplishes the processes for relative protein quantification, along with interactive visualization modules, including the novel VIQoR plot. We implemented two parsimonious algorithms to solve the protein inference problem, while protein summarization is facilitated by a well established factor analysis algorithm called fast-FARMS followed by a weighted average summarization function that minimizes the effect of missing values. In addition, summarization is optimized by the so-called Global Correlation Indicator (GCI). We test the tool on three publicly available ground truth datasets and demonstrate the ability of the protein inference algorithms to handle degenerate peptides. We furthermore show that GCI increases the accuracy of the quantitative analysis in data sets with replicated design.

**Availability and implementation:** VIQoR is accessible at: http://computproteomics.bmb.sdu.dk:8192/app_direct/VIQoR/

The source code is available at: https://bitbucket.org/vtsiamis/viqor/

**Supplementary information:** Supplementary data are available at *Bioinformatics* online.

## 1 Introduction

The last decades mass spectrometry (MS) has established as the preferred technology in quantitative proteomics (Gygi *et al*., 1999; Aebersold and Mann, 2003). High-throughput bottom-up analysis is the most used MS-based approach for the identification and quantification of thousands of proteins and the characterization of post-translational modifications (PTMs) in complex biological samples (Larsen *et al*., 2006; Zhang *et al*., 2017). Using this approach, the protein samples are enzymatically digested into shorter peptides to overcome problems with mass accuracy and complex MS/MS spectra. The resulting proteolytic peptide mixtures, either isotopically labeled (Thompson *et al*., 2003; Ross *et al*., 2004) or label-free (Cox *et al*., 2014), are typically first separated via liquid chromatography and then analyzed by tandem mass spectrometry (LC-MS/MS) to measure peptide mass, peptide intensity and the peptide fragment ions. Once peptides are identified and quantified using appropriate computational tools, the digestion needs to be computationally reverted by inferring proteins and summarize their abundance. These two tasks can be computationally challenging and require robust methods and algorithms that reduce the contributions of false peptide identifications and inaccurate abundance measurements to the quantified proteins (Huang *et al*., 2012).

Protein inference aims to determine the presence or absence of candidate proteins in a given sample, thus it can be considered as a protein identification process (Li and Radivojac, 2012). According to protein inference, a set of proteins presumed to be present in a sample is assembled from a set of identified peptides. In reality, however, solving the protein inference problem is not an easy task due to degenerate peptides, i.e. peptides that are shared by multiple proteins. To address this limitation, parsimony-based algorithms have been utilized to solve the protein inference problem (Yang *et al*., 2004; Alves *et al*., 2007; Zhang *et al*., 2007; Slotta *et al*., 2010; Koskinen *et al*., 2011; Uszkoreit *et al*., 2015). Algorithms that fall into this category rely on Occam’s razor principle, according to which, the most acceptable explanation of an occurrence is the simplest one. In this sense, the solution to the protein inference problem is a subset of the candidate proteins that explain the presence of all identified peptides in a sample in the most minimal way, therefore a solution to the NP-hard problem of minimum set cover (Karp, 1972; Huang *et al*., 2012; Li and Radivojac, 2012).

Parsimony-based algorithms are capable of creating minimal protein sets while handling degenerative peptides (Audain *et al*., 2017; The *et al*., 2018). However, they don’t utilize peptide quantitative measurements to infer proteins and are not providing estimations of protein concentrations. Simple methods that traditionally have been used to summarize protein concentrations, such as the average or median of the 3 most intense (top-3) or all peptide abundances have been shown to inherit the issues related to the current experimental and technical limitation of peptide quantification and identification (Goeminne *et al*., 2015). Recently, Diffacto, a tool for relative protein quantification has been developed to address these issues (Zhang *et al*., 2017). The tool applies the factor analysis method called fast-FARMS to extract the covariance between the observed peptide abundances for each identified protein. Based on that, individual weights are assigned to the peptides and in that way, unrepresentative or erroneous peptides can be eliminated. The protein concentrations are then calculated by a weighted average summarization of the remaining peptides. This approach has been demonstrated to increase the accuracy of protein abundance summarization in Diffacto. Nevertheless, the authors suggest an arbitrary peptide weight threshold of 0.5 to filter incoherent peptides, due to the bimodally distributed weights between 0 and 1. To date, a data-centric approach to obtain the optimal threshold and thus more accurate protein quantifications, is missing.

At the moment the available solutions that implement these processes are mostly command-line tools, scripts, libraries or are embedded in software suites(Breitwieser *et al*., 2011; Fischer and Renard, 2016; Zhang *et al*., 2017; Uszkoreit *et al*., 2019; Gatto *et al*., 2021). To our knowledge, there is still no tool for protein inference that provides an interactive visualization schema on the quantitative level of peptides, PTMs and proteins. Considering the superiority of advanced computational methods for protein inference and summarization and the importance of visualization tools as an essential step to understand and interpret proteomics data, we have developed VIQoR, a user-friendly web service for visually supervised protein inference and protein summarization. The tool accepts peptide or peptide-to-spectrum matches (PSMs) quantitative reports of any type of proteomics experiment (labeled or label-free) that involves at least four samples. VIQoR utilizes the Occam’s razor principle to infer proteins via two implemented parsimonious algorithms and adopts fast-FARMS to conduct a factor analysis that enables a weighted protein summarization. All processes are coupled to novel interactive visualization modules, user-friendly parameter optimization, feedback on data quality and configurable acquisition of high resolution graphs.

## 2 Materials and Methods

The current framework of VIQoR accomplishes all processes that follow peptide identification and are involved in protein quantification. The user can navigate through a user-friendly web interface and perform these processes along with interactive visualizations, as it is illustrated in the framework at Fig. 1. We present here the methods employed for protein inference and protein summarization. Additionally, the visualization methods implemented for VIQoR are listed and briefly explained.

**Fig. 1.**
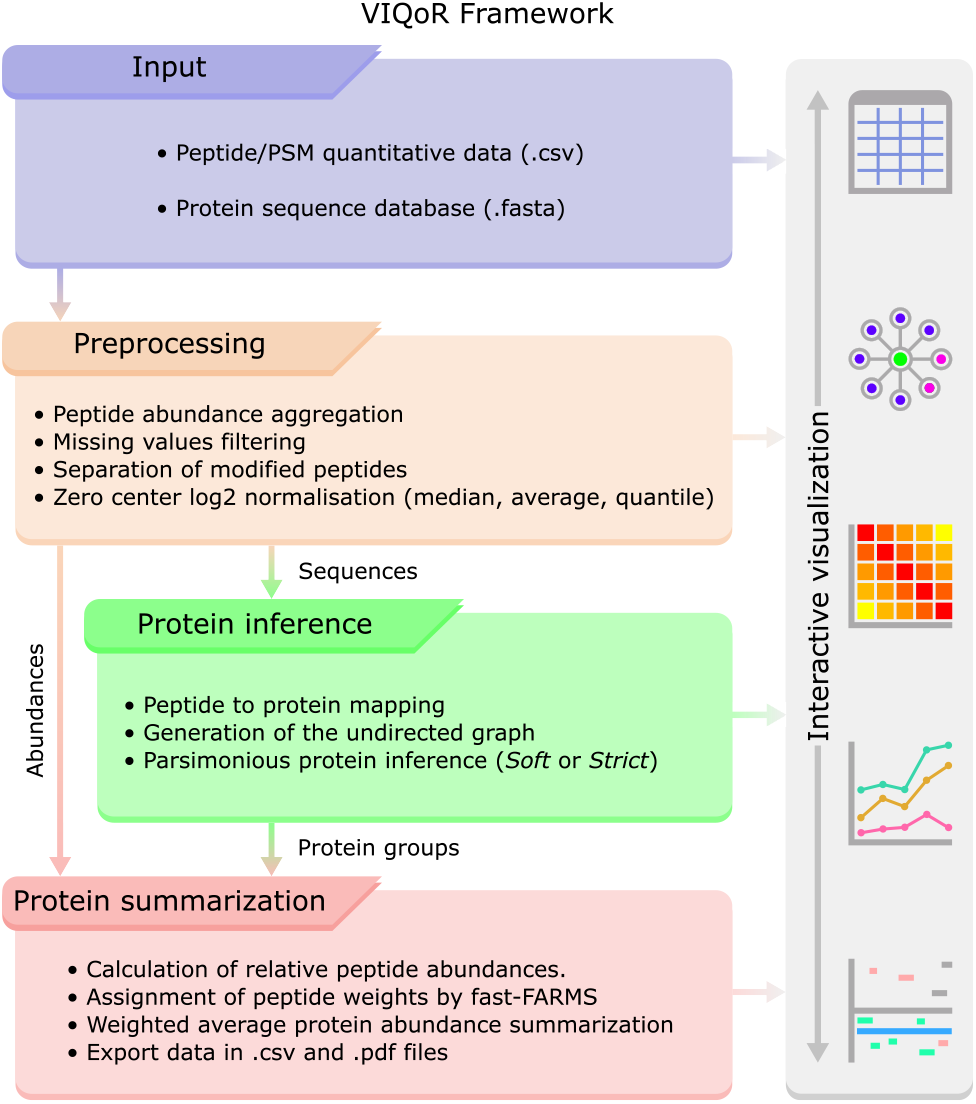
Schematic representation of VIQoR’s main framework: the peptide quantitative data and the protein sequence database files are imported in the Input tab (blue); the user can apply different filtering tasks and data transformations in the Preprocessing tab (orange); the sequence database and peptide sequences are used in the Protein Inference tab to create a minimal protein group set (green); protein quantification is performed in the Protein Summarization tab (red). The user can inspect and visualize the results of each process in the corresponding analysis tab interface, create new visualizations or access graphs generated in previous tabs to visually compare proteomics data of different levels.

### 2.1 Parsimony-based protein inference

Parsimonious algorithms model the relation between proteins and associated peptides on bipartite graphs and report a minimal set of proteins for each connected component. In graph theory, a connected component is a maximal subgraph in which every pair of vertices are connected by a path of edges. For each connected component, parsimonious algorithms typically report the proteins connected to the largest number of unexplained peptides, iteratively, until the presence of all peptides is justified (Huang *et al*., 2012; Li and Radivojac, 2012). Protein inference in VIQoR comprises two parsimonious algorithms that extend the classic parsimony by introducing additional criteria. These criteria enable the protein grouping. Therefore we will use the term ‘protein group’ to refer to the protein(s) inferred during each iteration and reported in the protein group set *P_Reported_*.

Given a connected component of a peptide set *p* related to a set of proteins *P*.

#### Strict parsimony

1. Append to set *P_Reported_* the protein *P_Leading_* of set *P* that is connected to the most peptides of set *p*. In case of multiple proteins *P_Leading_* with identical peptide sets *p_Leading_* they form a protein group (1st Criterion).
2. Report along with the appended protein group the associated peptide set *p_Leading_*.
3. Remove the peptide set *p_Leading_* from set *p*.

Continue iteratively the above steps until the set *p* is empty, and thus the presence of all identified peptides is explained uniquely by the minimal protein group set *P_Reported_*, a subset of *P*.

#### Soft parsimony

The peptide set *p_Initial_* denotes a copy of the initial peptide set *p*.

1. Append to set *P_Reported_* the protein *P_Leading_* of set *P* that is connected to the most peptides of set *p*. For all the remaining proteins in set *P*, in case their associated peptides in *p_Initial_* are a subset of *p_Leading_*, they form a protein group with *P_Leading_* (2nd Criterion).
2. Report along with the appended protein group the associated peptides of *P_Leading_* in set *p_Initial_* (3rd Criterion).
3. Remove the peptide set *p_Leading_* from set *p*.

Continue iteratively the above steps until set *p* is empty, and thus the presence of all identified peptides is explained at least once by the minimal protein group set *P_Reported_*, a subset of *P*.

Moreover, for each connected component VIQoR can report a maximal protein group set as a direct peptide to protein pairing, without employing the principle of parsimony. In this case, the protein group set *P_Reported_* is equal to set *P*. An example of each approach as a comparison applied on the same connected component is illustrated in Supplementary Fig. S2.

### 2.2 Relative protein abundance summarization

Protein abundance summarization in VIQoR is facilitated by a Bayesian factor analysis method called fast-FARMS, which is a reimplementation of FARMS (Factor Analysis for Robust Microarray Summarization), an algorithm originally developed to summarize probe level oligonucleotide microarray data based on their extracted covariance (Hochreiter *et al*., 2006). Fast-FARMS has been implemented for Diffacto (Zhang *et al*., 2017) to estimate relative protein abundances and has been previously employed by ComplexBrowser (Michalak *et al*., 2019) and CoExpresso (Chalabi *et al*., 2019) for protein complexes quantification. It has been demonstrated that fast-FARMS is a robust method that can serve efficiently as an additional post-identification filtering process on abundances of peptides that appear to derive from the same proteins. Consecutively, it has been shown that it outperforms in accuracy other popular protein abundance summarization approaches (Zhang *et al*., 2017).

The method assumes that the real protein concentration ratio is proportional to the abundance ratios of the proteolytic peptides, therefore models their relation linearly with an additional Gaussian noise. The following steps are performed for each inferred protein group:

1. Calculate the relative abundances of the log2 transformed and zero-centered normalized peptides.
2. Apply the factor analysis of fast-FARMS to assign individual peptide weights.
3. Apply minimum weight threshold and summarize protein concentration ratios as the weighted average of the remaining representative peptides.

The complete factor analysis algorithm is described in FARMS manuscript (Hochreiter *et al*., 2006), while its application in proteomics is presented by Diffacto authors (Zhang *et al*., 2017).

### 2.3 VIQoR plot

VIQoR plot is an interactive visualization module that combines both quantitative and amino acid sequence information of a selected protein between two samples or conditions. The horizontal axis denotes the amino acid position on the selected protein sequence and the vertical axis corresponds to the log2 fold change of the protein and peptide abundances between the two selected samples or conditions. The peptide sequences (modified and unmodified) that correspond to the selected protein are visualized as horizontal bars with respect to their position in the protein sequence. Additionally, the individual PTMs are mapped accordingly and are annotated on the bars of the modified peptides.

### 2.4 Global correlation index

The protein summarization in VIQoR employs the weights assigned to the peptides to assess the peptide coherence for the protein they identify. Hence, not only the weights can lower the effect of erroneous measurements but an additional application of a minimum threshold can eliminate their contribution. Consequently, in VIQoR the user can select a suitable cut-off weight value. In replicated experiments, the Global Correlation Index (GCI) is a line plot that summarizes the correlation of protein expression of replicates within the same conditions for a set of predefined peptide weight thresholds. Assuming a replicated experiment of *CR* samples comprising *C* (*C* > 1) conditions and *R* (*R* > 1) replicates, the protein expression matrix of all quantified protein groups is calculated for each peptide weight threshold *w* = 0, 0.1, 0.2, …, 1. Then, for each threshold *w*, we calculate the *Within* correlation *GCI*(*w*) as the mean of all pairwise *CR*(*R* − 1)/2 Pearson correlation coefficients between the samples of the same condition. Correlations between identical samples are excluded.

### 2.5 Software implementation

VIQoR is implemented in R (version≥4.0.0) and customized by JavaScript. For the user interface, the R packages Shiny (version≥1.4.0), shinydashboard (version≥0.7.1), DT (version≥0.13), shinyjs (version≥2.0.0), shinyBS (version≥0.61) and shinycssloaders (version≥0.3) have been used. The visualization functionality of the tool is facilitated by Plotly (version≥4.9.2), heatmaply (version≥1.1.0) (Galili *et al*., 2018), networkD3 (version≥0.4) and webshot2 (version≥0.0.0.9000). Data analysis modules were developed by the R packages igraph (version≥1.2.5), biostrings (version≥2.56.0), protr (version≥1.6.2) (Xiao *et al*., 2015), dplyr (version≥1.0.3), preprocessCore (version≥1.50.0). Additionally, PeptideMapper from the compomics-utilities library (version≥4.12.9) (Barsnes *et al*., 2011) is used for fast peptide to protein sequence mapping (Kopczynski *et al*., 2017).

VIQoR can be run independently on a local shiny server, via RStudio, directly deployed using its docker container or accessed as a web service at http://computproteomics.bmb.sdu.dk:8192/app_direct/VIQoR/.

### 2.6 Ground truth datasets

#### Hybrid dataset 1

20 tryptic peptide mixtures of human and yeast cell lysates and bovine serum albumin (BSA) of known concentrations were analyzed using label-free data-dependent acquisition (DIA) mass spectrometry on an Orbitrap Q-Exactive Plus coupled to an LC system (Thermo Fisher Scientific), in three technical replicates. The amount of the human peptides among the conditions is reduced linearly (from 1000 to 1 ng), the amount of BSA peptides is increased exponentially (from 0.02 to 120 ng), while the proportion of yeast peptides added maintain the total peptide amount in all samples at 1001 ng. An additional sample was analyzed to serve as an internal standard with 350 ng, 7.6 ng and 636 ng of human, BSA and yeast peptides respectively. Morpheus (revision 165) was used for peptide identification, with a q-value threshold of 0.01 and peptides were quantified by DeMix-Q workflow (Zhang *et al*., 2017).

#### Hybrid dataset 2

Two hybrid samples consisting of human, yeast and E. coli proteins in known concentration ratios of 1:1, 10:1 and 1:10, respectively, were digested by trypsin. Three technical replicates of proteolytic peptide mixtures for each sample were separated by an HPLC system (Eksigent) and followed by label-free data-dependent acquisition (DDA) analysis on a 6600 TripleTOF (ABSciex) mass spectrometer. Acquired MS/MS spectra were identified by PLGS at 1% FDR and peptide abundances reported by ISOQuant. *Hybrid dataset 2* is provided by the LFQbench R package under the name *hye110* (Navarro *et al*., 2016).

#### Spike-in dataset

tryptic peptides of mouse and rat lyophilized CERU in four known concentration ratios were spiked into a background mixture of trypsin digested albumin and IgC depleted human plasma proteins, for an iTRAQ 4-plex designed experiment. Rat (P13635) and mouse (Q61147) CERU concentration ratios are 1:2:4:10 and 10:5:2:1 respectively, while human protein concentrations remained stable (1:1:1:1). The labeled sample was analyzed on a hybrid LTQ-Orbitrap XL mass spectrometer (Thermo Fisher Scientific) coupled to HPLC nanoflow system (Agilent). Mascot (version 2.3) and Phenyx (version 2.6.1) conducted peptide identification with individual peptide identification false positive rate lower than 0.1%. The R package Isobar offers the *Spike-in dataset* under the name *ibspiked_set1* (Breitwieser *et al*., 2011).

### 2.7 Quantification benchmarking

The peptide abundances for *Spike-in dataset* were calculated as the sum of their PSM intensities in each sample. For all three datasets the peptides quantified in at least 2 samples were taken, log2 transformed and zero-center normalized by median. Peptide abundances were relatively expressed to the average abundance of each peptide in all samples. Factor analysis hyperparameters weight and mu were both set to 0.1.

For the *Hybrid dataset 1*, Pearson correlation coefficient was used to investigate the relation between the protein group concentration profiles and the 21 actual concentrations. Average correlation is then calculated as the arithmetic mean.

The performance of the protein quantification in the *Hybrid dataset 2* was assessed with a Receiver Operating Characteristic (ROC) curve. True positives (TP) are the quantified protein groups with expression levels falling within the increasing but not overlapping cut-off ranges around the real concentrations and with at least one protein member of the corresponding species.

To assess the accuracy of quantification for *Spike-in dataset* the Standard Error of the Estimate (SEE) was calculated between the real and the estimated concentrations.

## 3 Results

VIQoR is a user-friendly R-based web application that facilitates protein inference, protein abundance summarization and proteomics data visualization. The flowchart of the implemented framework is illustrated by Supplementary Fig. S1. An extensive description of the input data requirements and the tool’s functionalities can be found in the manual (Sup. File S1). In this study we first examine the two implemented parsimonious approaches and demonstrate their ability to infer and separate proteins with largely similar sequences. Second, we demonstrate how the GCI values can be used to optimize the accuracy of quantification by fast-FARMS and lastly, we provide use case scenarios regarding the application, visualization and interpretation of VIQoR plots.

### 3.1 Parsimony criteria and homologous protein separation

The two implemented parsimonious algorithms add flexibility to the classic principle of parsimony to handle degenerate peptides by introducing additional criteria. The *Strict* parsimony groups only proteins that share exactly the same peptides, while *Soft* parsimony enables the merging of overlapping proteins and maintains a copy of the initial peptide set until convergence. Consequently, by the *Soft* approach all peptides are explained at least once, unlike the *Strict* approach where all peptides are explained only once. In other words, the two algorithms differ by criteria of ‘strictness’ applied when addressing degenerate peptides. The third approach of no parsimony then corresponds to the ‘softest’ side of that spectrum as the direct peptide to protein pairing. A practical example of the three approaches on the same connected component is illustrated in Supplementary Fig. S2. When *Soft* parsimony is applied (Supplementary Fig. S2C), the tolerance in accepting degenerate peptides is higher and protein grouping is promoted. On the other hand, *Strict* parsimony distinguishably separates and reports the most evident proteins until the presence of all peptides is uniquely explained (Supplementary Fig. S2B).

To demonstrate the differences of the inference approaches to more complex systems than a single connected component, we tested their performance on the two hybrid datasets. *Hybrid dataset 1* contains the intensities of 38.794 unique peptides and *Hybrid dataset 2* contains 13.444 measured peptide features, from which 12.667 are unique. The datasets fit ideally to this demonstration since they contain a large number of degenerate peptides, i.e. 1794 and 1150 individually. Both parsimonious approaches reported the same number of protein groups for the two datasets (including one-hit wonders), being 7118 and 3589, respectively (Supplementary Table S1). Non-parsimonious inference reported more protein groups compared to the parsimony driven methods. More specifically, 7263 groups for *Hybrid dataset 1* and 5016 for *Hybrid dataset 2*. For both datasets, the *Strict* algorithm reported the lowest number of protein groups identified by more than one peptide, while inference without parsimony reported the most, something that clearly derives from the different levels of tolerance addressed to degenerate peptides. Furthermore, we examined the overlap between the protein groups reported by the three approaches (Supplementary Fig. S3). Interestingly, in both datasets and for an overall comparison, the groups yielded *Soft* parsimony in terms of peptide composition are a subset of the groups reported when no parsimony was employed. Additionally, for larger protein groups in terms of protein composition, *Strict* appears to be a subset of the protein groups reported by the *Soft* approach.

To investigate further the parsimony criteria on proteins with prior knowledge of high amino acid sequence similarities, we used the *Spike-in dataset* since its spiked proteins are orthologs of known concentrations. In this comparison we exclude the non-parsimonious approach because unlike the two parsimonious algorithms, it distributes the shared peptides to all related proteins. The dataset is a PSM report consisting of 14991 PSMs of 1653 unique peptides. The samples of the *Spike-in dataset* contain ceruloplasmin orthologs from mouse (CERU_MOUSE), rat (CERU_RAT) and human (CERU_HUMAN). All three proteins follow the concentration levels of their respective species throughout the four iTRAQ channels. In total 101 peptides were identified for the three proteins and 20 of them are degenerate as shown in Fig. 2. The resulting quantitative protein profiles for CERU_HUMAN and CERU_MOUSE according to the two parsimonious approaches are illustrated in Fig. 3. The *Soft* approach (Fig. 3A,C) added 8 and 18 degenerate peptides (red traces) to CERU_HUMAN and CERU_MOUSE, respectively. The majority of these peptides do not follow the real concentrations of the two species. *Strict* parsimony (Fig. 3B,D) appears to separate the shared peptides in a more robust way and reduce the variance of the peptide abundances in every sample. The erroneous assignment of peptides in the *Soft* approach affects the estimation of protein concentration by fast-FARMS. The summarized protein abundance for both proteins with *Strict* parsimony is following more accurately the real concentrations (Fig. 3, black dashed lines) compared to *Soft*, which is confirmed by the SEE scores. The estimated error is lower when the two proteins are inferred by the *Strict* approach, 0.125 and 0.263 for CERU_HUMAN and CERU_MOUSE respectively.

**Fig. 2.**
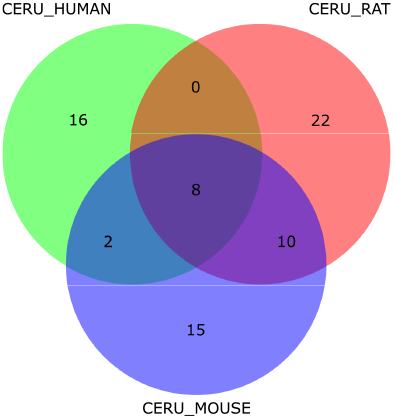
Ceruplasmin proteins of the considered species in exhibit different sets of peptides. Venn diagram of the peptide composition for the three ceruplasmin proteins in the Spike-in dataset.

**Fig. 3.**
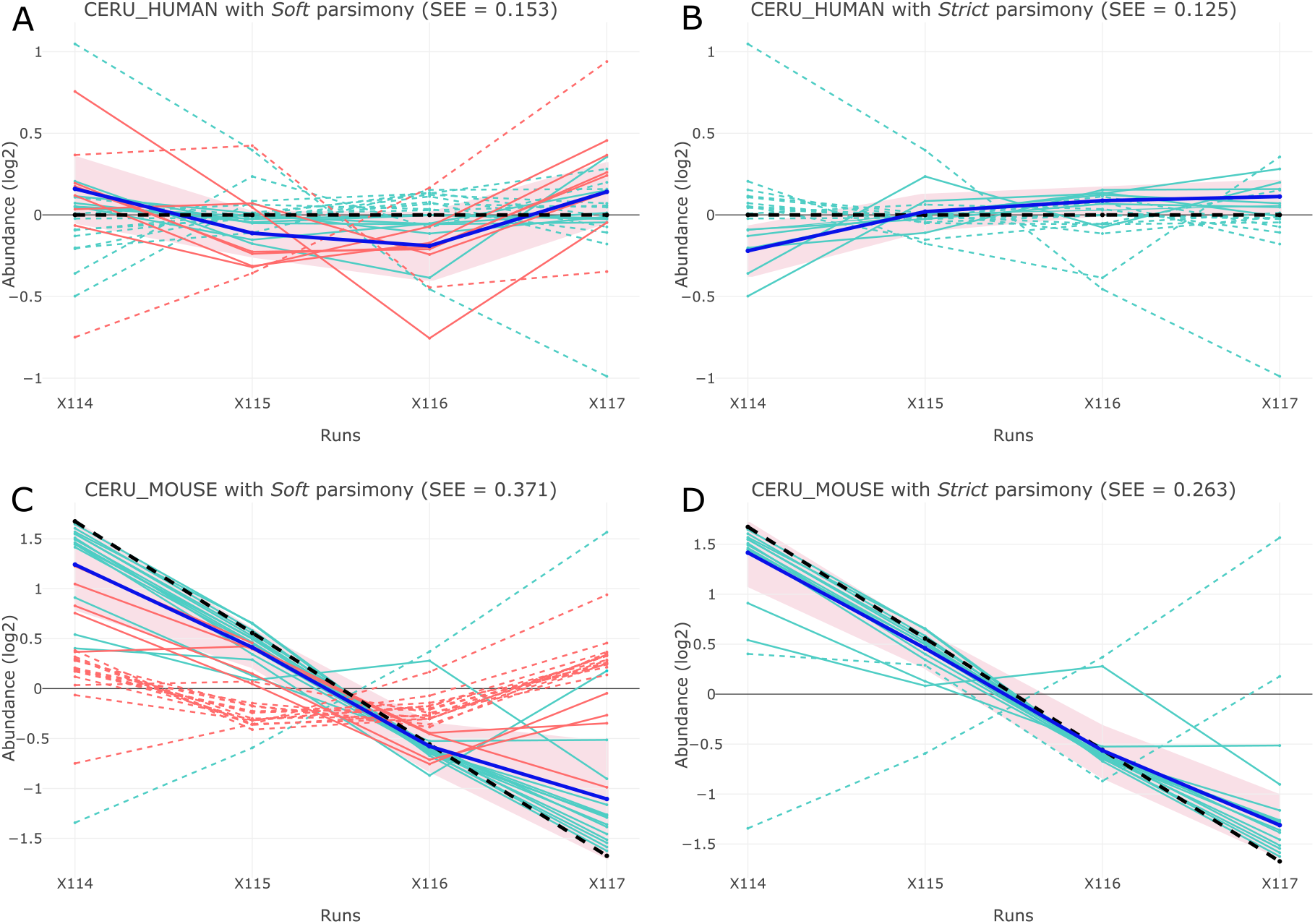
Strict parsimony increases the quantification accuracy in ortholog proteins. Protein inference with Soft and Strict parsimony for spiked in proteins CERU_HUMAN and CERU_MOUSE. A-D – Protein expression (thick blue) and peptide abundance (cyan and red) line plots of CERU_HUMAN and CERU_MOUSE proteins. Protein expression is summarized by fast-FARMS with a peptide weight threshold of 0.5. Solid lines correspond to peptides of weight ≥ 0.5 and dashed lines to the eliminated peptides of weight < 0.5. Peptides assigned to the same protein group by both parsimony methods are colored cyan and red otherwise. Pink highlighted area corresponds to the interval of ± one standard deviation. The SEE statistics are calculated as a comparison to the standardized real concentration of the proteins (thick dashed lines). A – CERU_HUMAN summarization by Soft parsimony. B – CERU_HUMAN summarization by Strict parsimony. C – CERU_MOUSE summarization by Soft parsimony. D – CERU_MOUSE summarization by Strict parsimony.

### 3.2 Similarity within samples of the same experimental condition as indicator of quantification quality

Protein quantification utilizing fast-FARMS in Diffacto have been reported not only to summarize in total more protein groups but also to surpass other summarization techniques in accuracy and precision (Zhang *et al*., 2017). We analyzed the two hybrid datasets with VIQoR to verify the robustness that fast-FARMS can provide in protein summarization and furthermore to present a detailed comparison of the parsimonious approaches. For *Hybrid dataset 1*, regardless of the protein inference approach and peptide weight threshold, the average correlation of protein abundances for the 21 real concentrations is higher than 0.77 (Fig. 4A, dashed lines). Similarly, the evaluation of protein summarization of *Hybrid dataset 2* resulted in AUC scores higher than 0.92 in all cases (Fig. 4B, dashed lines). For both datasets, the *Strict* parsimony improved the summarization by more accurately reproducing the known protein changes. Despite the generally accurate performance of the factor analysis in the two datasets, the different elimination of incoherent peptides by applying different weight thresholds has a considerable impact on the accuracy of protein quantification. VIQoR employs the expected higher similarity between sample replicates to estimate the most appropriate value for the weight threshold. For that purpose, the GCI module in VIQoR’s user interface presents the *Within* correlation traces. These are illustrated as solid lines in Fig. 4.

**Fig. 4.**
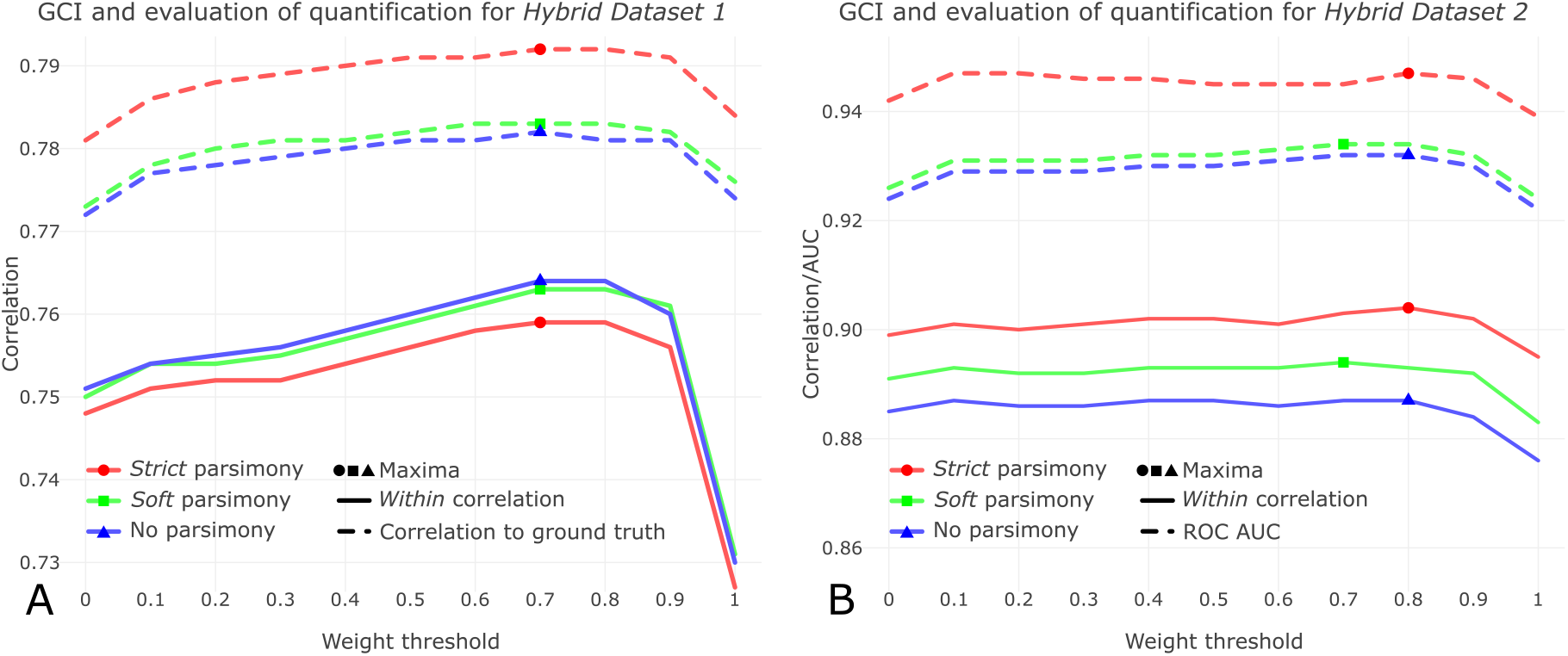
GCI optimizes the performance of protein summarization by fast-FARMS. GCI scores and the evaluation of protein quantification per peptide weight threshold. Solid lines correspond to the Within correlation scores between sample replicates, as reported by GCI module. Dashed lines show the evaluation of quantification and denote either the summarized correlation of protein expression to real concentrations for Hybrid dataset 1 (A) or the AUC scores for Hybrid dataset 2 (B). Red lines represent the quantitative analysis followed by protein inference with Strict parsimony, Green with Soft parsimony and Blue without parsimony. In all six evaluations (three for each dataset), the peptide threshold with the optimum quantification output corresponds to the weight that maximizes the GCI.

As expected, the similarity in protein expression within each condition results in high scores in GCI (Fig. 4). The *Within* correlation of GCI for *Hybrid dataset 1* is higher than 0.72 regardless of the weight threshold value (Fig. 4A, solid lines). The GCI is maximized for all inference approaches at the weight threshold of 0.7, with scores of 0.759, 0.763 and 0.74 for *Strict*, *Soft* and without parsimony, respectively. Similarly, the *Within* correlation is higher than 0.87 for all instances for *Hybrid dataset 2* (Fig. 4B, solid lines). For this dataset, the GCI has maximum peaks in different weight thresholds, with correlation of 0.894 at weight of 0.8 for *Soft* parsimony and correlations of 0.904 and 0.887 for *Strict* and without parsimony summarization at weight of 0.7. The maximum peaks of GCI for all evaluations align perfectly to the peptide weight thresholds that result in the optimal protein quantification (Fig. 4). This shows that the GCI allows optimizing the value of the weight threshold and subsequently increases the accuracy and precision of protein quantification in VIQoR.

Nevertheless, this metric does not necessarily determine the most suitable protein inference approach. For instance, the approach that has the highest GCI scores for *Hybrid dataset 1* is the protein inference without parsimony, followed by the *Soft* and *Strict* approaches. However, as stated above, the *Strict* parsimony provides the optimal quantification for that dataset.

### 3.3 Protein quantitative analysis with simultaneous visual inspection

In VIQoR the processes of protein inference and protein summarization are coupled to visualization modules for the inspection of specific results regarding protein groups and samples. The tool provides interactive tables to display quantitative data, connected component graphs of the inferred protein groups, Pearson correlation heatmaps of the protein and peptide abundances and peptide-centric line plots of protein group profiles. All the generated visualizations can be exported as vector graphics in .pdf format.

In order to visualize (modified and unmodified) peptide and protein quantifications together with the sequence coverage of the selected quantified protein groups, we implemented the VIQoR plot (Fig. 5). This plot shows log2 fold changes of the summarized protein expressions and the normalized peptide abundances between two samples or conditions. The peptides are arranged horizontally according to their position in the protein sequence, while the vertical distance equals the corresponding log2 fold change. In addition to the peptides involved in the summarization process, the module also visualizes the incoherent peptides eliminated by the peptide weight threshold (grey color). The VIQoR plot provides supplementary information via tooltips, such as amino acid sequence and its length, the position of the peptide on the reference sequence, the log2 fold change and the peptide weight value assigned by fast-FARMS.

**Fig. 5.**
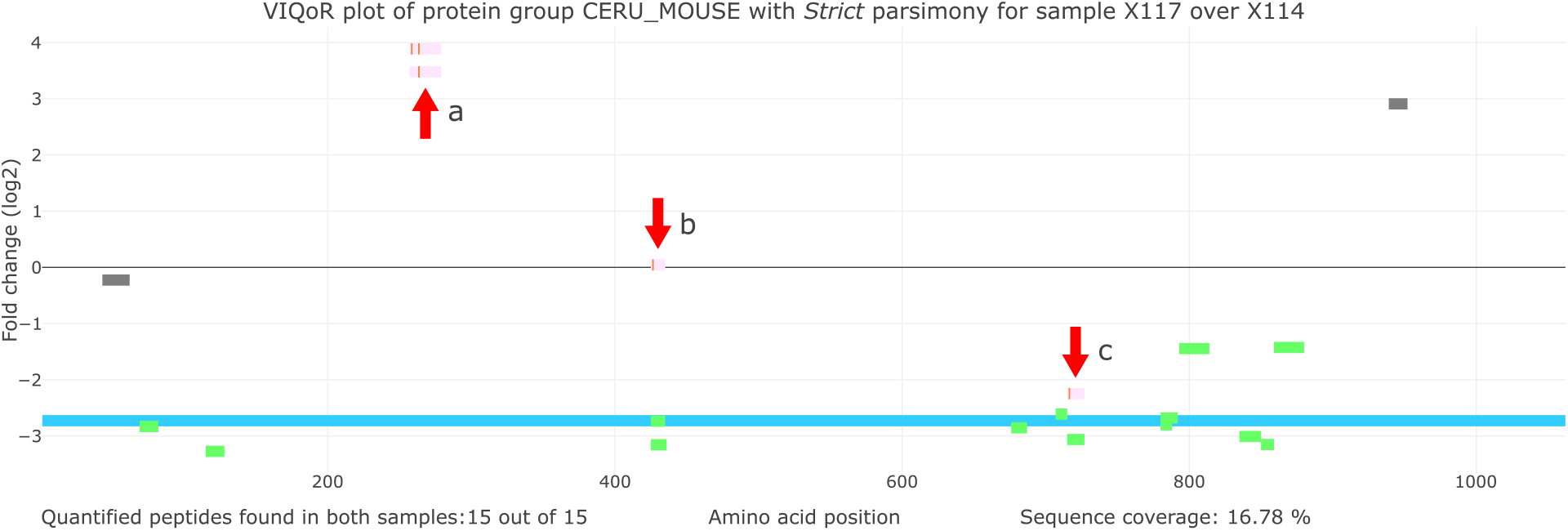
Visual exploration of peptide protein coverage that includes localization and quantification of post translationally modified peptides. VIQoR plot for CERU_HUMAN of the Spike-in dataset. The protein is inferred with Strict parsimony. Blue stripe corresponds to the protein, green segments to the peptides of weight ≥ 0.5, grey to the eliminated peptides and pink to the modified peptides. Orange annotations denote the modification type and site.; a - Modified peptides with phosphorylation sites Y264 and Y259, Y264, respectively.; b - Modified peptide with phosphorylation site Y427.; c - Modified peptide with phosphorylation site F717

Fig. 5, demonstrates a VIQoR plot of the CERU_MOUSE of the *Spike-in dataset* inferred by *Strict* parsimony between the first and fourth iTRAQ channels. The log2 fold change of the protein expression is −2.72, which corresponds to a concentration ratio of 0.15, therefore reasonably close to the real ratio of 0.1. The 13 peptides with assigned weight over 0.5 are colored green. The two eliminated peptides in grey color are the same incoherent degenerate peptides with dotted traces in Fig. 3D. Generally, VIQoR plot shows how an optimal peptide threshold reduces the overall variance within all the considered peptides.

VIQoR provides flexible integration of modified peptides in the analysis. Our tool supports peptide or PSM reports of both modified and unmodified peptides. During the data preprocessing the modified peptides are separated by modification type and the user can select which PTMs will be proceeded for visualization purposes (Fig. 1). The abundance of the selected modified peptides are not considered in protein summarization. However, the peptides with these PTMs are still mapped on the inferred protein sequences and used in the VIQoR plot. The PTM abundances are not expressed relatively to the protein concentration but as the log2 fold change of the modified peptide intensities. In that way, the PTM expression patterns can be projected on the protein expression profile.

The changes of the modified peptides abundances do not necessarily suggest changes of PTM levels relative to the protein, yet the protein expression level should be considered too. Therefore, multiple possible combinatorial scenarios between modified peptides and protein abundances arise (Kim *et al*., 2016). To present some of the different usage scenarios of the VIQoR plot regarding the behavior of the modified peptides, we retrieved 4 phosphorylated peptides of CERU_MOUSE from PhosphoSitePlus (Hornbeck *et al*., 2015), that contain the phosphorylation sites F717, Y427, Y264 and Y259, Y264. The modified peptides were assigned to *Spike-in dataset* measurements that follow the protein concentrations of mouse, human and rat, respectively. We then added the peptides to the *Spike-in dataset* (cases *a* − *c* in Fig. 5). In case *a*, the modified peptides follow opposite regulatory behavior to the one of the protein. A second scenario is shown in case *b*, where the modified peptide abundance remains unchanged and the protein expression is down-regulated. This could correspond to a proteoform of the selected protein that does not get altered. Finally in case *c*, the modified peptide and the protein follow the same changes and thus are co-regulated.

The VIQoR plot simplifies understanding the full behavior of a protein including biologically relevant relative changes of its modified peptides that do not follow the common trend of the unmodified protein. Additionally, since each modification type is annotated in a different color, the user can obtain thorough insight into the relation between the expression of different modification types.

## 4 Discussion

In this study, we present VIQoR, a user-friendly online tool that processes quantitative PSM/peptide reports and performs protein inference, relative protein abundance summarization and simultaneous visualization. The imported reports can derive from any type of experiment or MS analysis. Hence, to evaluate the performance VIQoR we selected labeled and label-free ground truth datasets of both DIA and DDA studies.

We tested two parsimonious approaches for their performance with respect to degenerate peptides. *Strict* parsimony showed more robust separation of the degenerate peptides of ortholog proteins when compared to the *Soft* approach and also provided more accurate protein quantification, regardless of the peptide weight threshold. We also could show that the parsimony principle outperformed the standard method of direct protein assembly. However, our comparison of protein inference approaches was restricted to multi-species datasets that contain a large fraction of degenerate peptides and thus is not necessarily applicable to single species experiments. Therefore, *Soft* parsimony might be more appropriate for the analysis of other types of datasets, e.g. single species experiments. However, this will require further investigation.

Pearson correlation coefficients are often used to measure the variability and reproducibility of both gene expression microarray (Kuo *et al*., 2006) and quantitative proteomics (Perrin *et al*., 2013) experiments. We have applied the same principle to access the similarities of protein expression profiles between replicates. With that, we furthermore demonstrate that the maximization of the *Within* correlation in GCI increases the quantification quality. Indeed, for both ground-truth hybrid datasets, the most accurate protein quantification was achieved by the peptide weight threshold value that maximizes the GCI. We assume that a similar unsupervised approach for parameter optimization could fit in other methods involved in proteomics data analysis. However, as it is demonstrated by the analysis of *Hybrid dataset 1*, GCI scores are independent of the inference method and therefore they cannot indicate the most accurate approach. We suspect that this is an effect of the increased amount of shared peptides between the reported protein groups inferred by the non-parsimonious and *Soft approaches*, corresponding to 4.6% and 3.6% of the peptides respectively. We assume this higher number of shared peptides in turn increases the similarities among the summarized protein abundances.

Lastly, we have developed the VIQoR plot, a novel visualization that combines quantitative details about peptide and protein changes with amino acid sequence data. The plot can be used to compare the response of a protein group between two biological states and at the same time illustrates the quantitative patterns of all measured modified and non-modified peptides. This makes VIQoR valuable to PTMs analysis as it allows detailed views or PTM behavior with respect to the general trend of a protein. VIQoR plots along with the other interactive visualization modules can serve as an intuitive tool for exploratory analysis and data interpretation, while the user can browse and visualize data on peptide, PTM, protein and protein group level.

## Acknowledgements

V.T. This work would not have been possible without the useful discussions with Marija Nišavić and Maša Babović.

