## Supplementary material for "VIQoR: a web service for Visually supervised protein Inference and protein Quantification"

April 2021

### 1 Supplementary Figure 1

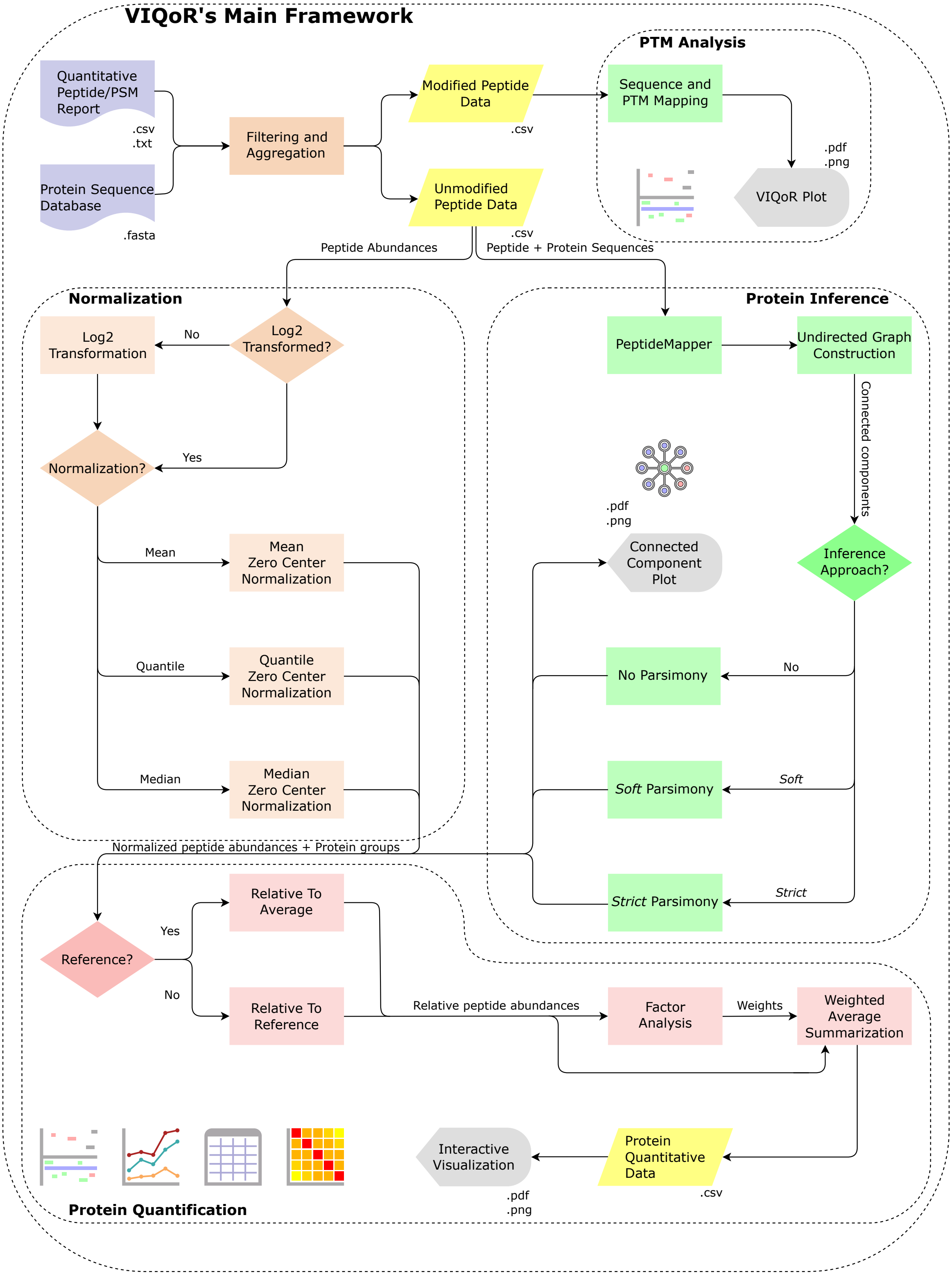


**Supplementary Fig. S1:** Flowchart of VIQoR's framework. Document symbols denote the input files and parallelograms the main input/output data types. Rectangles correspond to the performed processes, while diamond symbols to software interactions with the user. Display symbol is used for the visualization processes.

### 2 Supplementary Figure 2

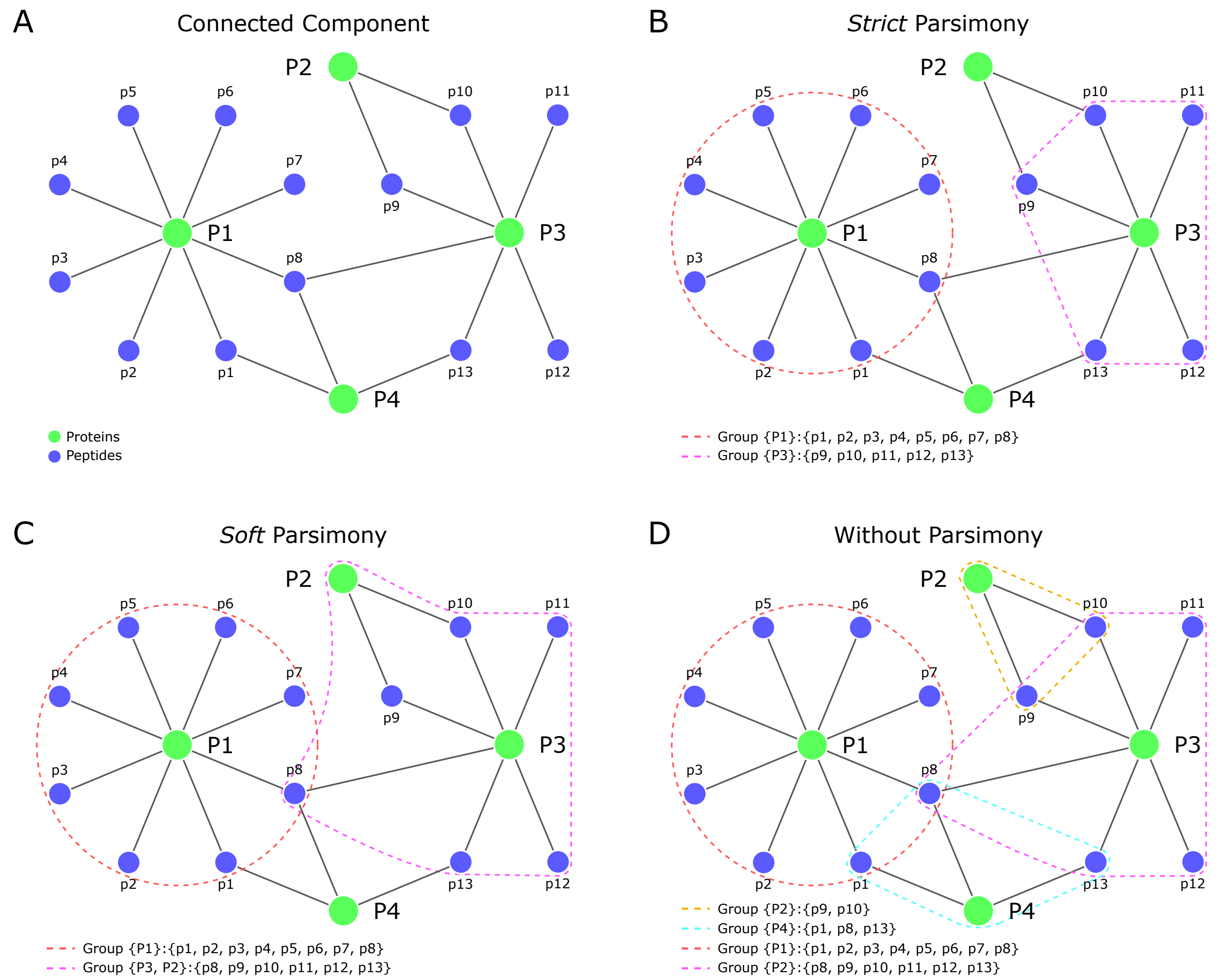


**Supplementary Fig. S2:** Example of three protein inference approaches applied on the same connected component. A - A connected component of a bipartite graph for peptide set $p = p1, p2, p3 \dots p12$ and protein set $P = P1, P2, P3, P4$. Green and blue nodes correspond to proteins and peptides respectively, while edges express their relation.; B - D - Dashed line contours specify the inferred protein groups.; B - Protein inference by *Strict* parsimony. $P1$ is the first $P_{Leading}$ protein connected to 8 peptides and $P3$ is the second $P_{Leading}$ protein connected to 5 peptides. The minimal protein set that explains the presence of all peptides in set $p$ is $P_{Reported} = P1, P3$; C - Protein inference by *Soft* parsimony with $P_{Reported} = P1, (P3, P2)$. Protein $P2$ forms a protein group with $P3$ due to the $2^n d$ Criterion. Peptide $p8$ is shared by both protein groups due to the $3^r d$ Criterion.; D - Direct peptide to protein pairing, without the involvement of parsimony.

### 3 Supplementary Table 1

| Inference | *Hybrid Dataset 1* | | *Hybrid Dataset 2* | |
|---|---|---|---|---|
| Method | All protein groups | > 2 peptide groups | All protein groups | > 2 peptide groups |
| *Strict* Parsimony | 7118 | 3838 | 3589 | 1365 |
| *Soft* Parsimony | 7118 | 4005 | 3589 | 1505 |
| No Parsimony | 7263 | 4063 | 5016 | 1644 |

**Supplementary Table S1:** Number of protein groups inferred by three different approaches for *Hybrid dataset 1* and *Hybrid dataset 2*. Columns with header "All protein groups" correspond to the total number of reported protein groups including those identified by a single peptide (one-hit wonders).

### 4 Supplementary Figure 3

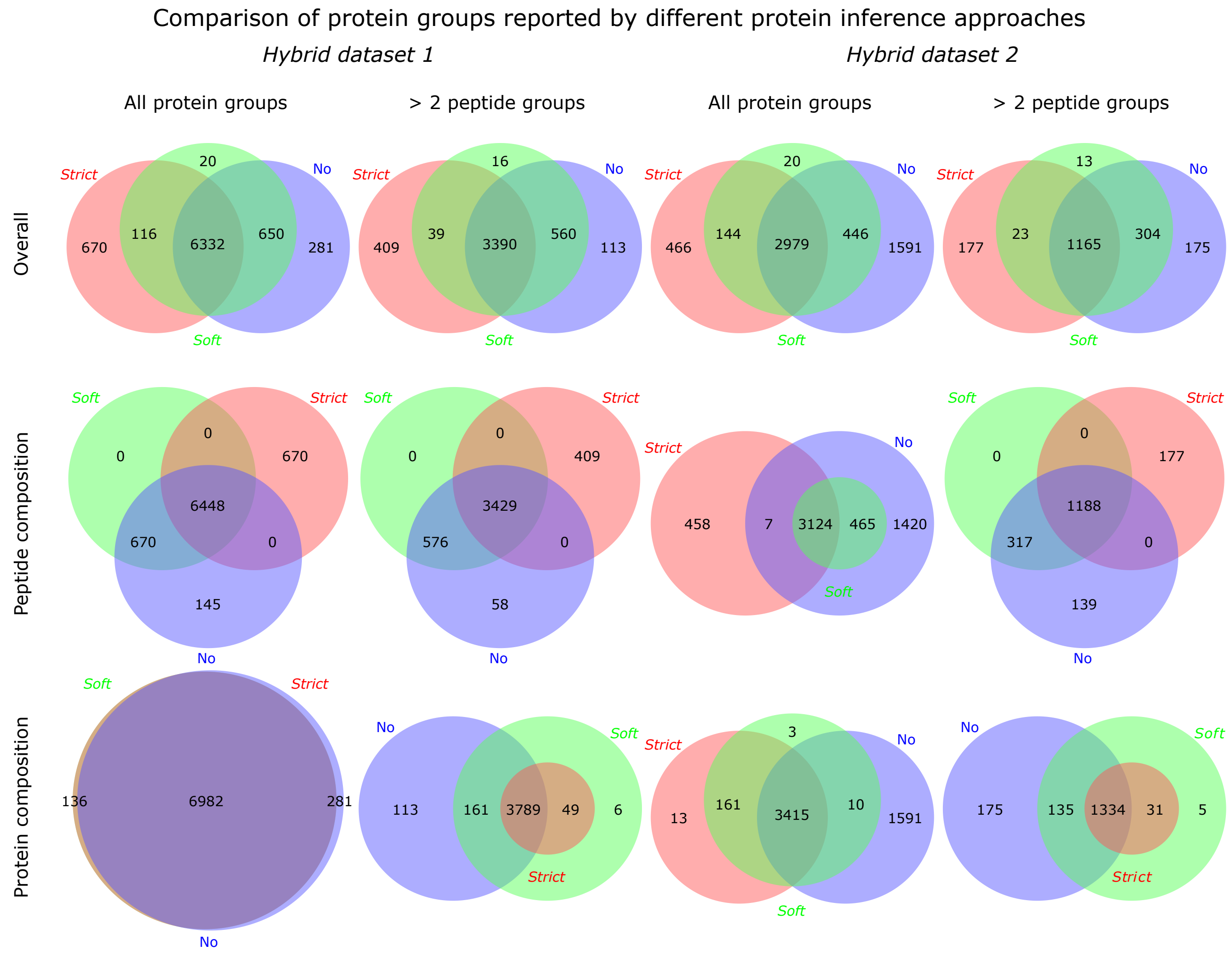


**Supplementary Fig. S3:** Comparison of protein groups reported by different protein inference approaches for *Hybrid dataset 1* and *Hybrid dataset 2*. Venn diagrams of overlapping protein groups in an overall, and peptide and protein composition comparison. Peptide composition refers to the set of peptides assigned to a protein group. Protein composition refers to the set of protein identifiers assigned to a protein group. The protein composition of two groups is equal if the union of their protein identifiers is identical to the composition of the group with the most proteins. Overall comparison considers both protein and peptide composition to the comparison.
